# Expanded GGGGCC repeat transcription is mediated by the PAF1 complex in *C9or72*-associated FTD

**DOI:** 10.1101/319376

**Authors:** Lindsey D. Goodman, Mercedes Prudencio, Luis F. Martinez-Ramirez, Matthews Lan, Michael Parisi, Leonard Petrucelli, Nancy M. Bonini

## Abstract

Expression of an expanded (G4C2)30+ repeat found in *C9orf72* is the most prominent mutation in familial FTD and ALS. An unbiased RNAi-based, large-scale screen in (G4C2)49-expressing *Drosophila* identified the CDC73/PAF1 complex (PAF1C) as a novel suppressor of (G4C2)49-toxicity. Downregulation of PAF1C, an activator of elongating RNAPII, caused suppression by reducing (G4C2)49-RNA levels. Remarkably, only PAF1C components Paf1 and Leo1 were selective for transcription of a long repeat expansion; transcript levels produced from shorter and longer repeat-containing transgenes were similarly affected by other components and Spt4, a previously identified transcriptional regulator of G4C2-repeats. Congruent with our fly data, *PAF1* and *LEO1* were upregulated in the frontal cortex of C9+ FTD patients and their expression correlated to expression of repeat-containing *C9orf72*. Surprisingly, this affect was specific to C9+ FTD versus C9+ ALS. This is the first evidence that PAF1C is playing a role in *C9orf72-*associated FTD. Further, PAF1C may affect other repeat-associated diseases.

**HIGHLIGHTS:**

- (G4C2)49-toxicity modifier screen highlights suppressors as RNAPII-transcription regulators
- Further insights into repeat mediated transcription by *Spt4/DSIF* in C9+ FTD/ALS
- Addition of PAF1C as an elongation complex important for promoting RNAPII-transcription of G4C2 repeats

- (G4C2)30+ transcription is mediated by Leo1 and Paf1 of the Paf1C complex, not by DSIF

## MODEL

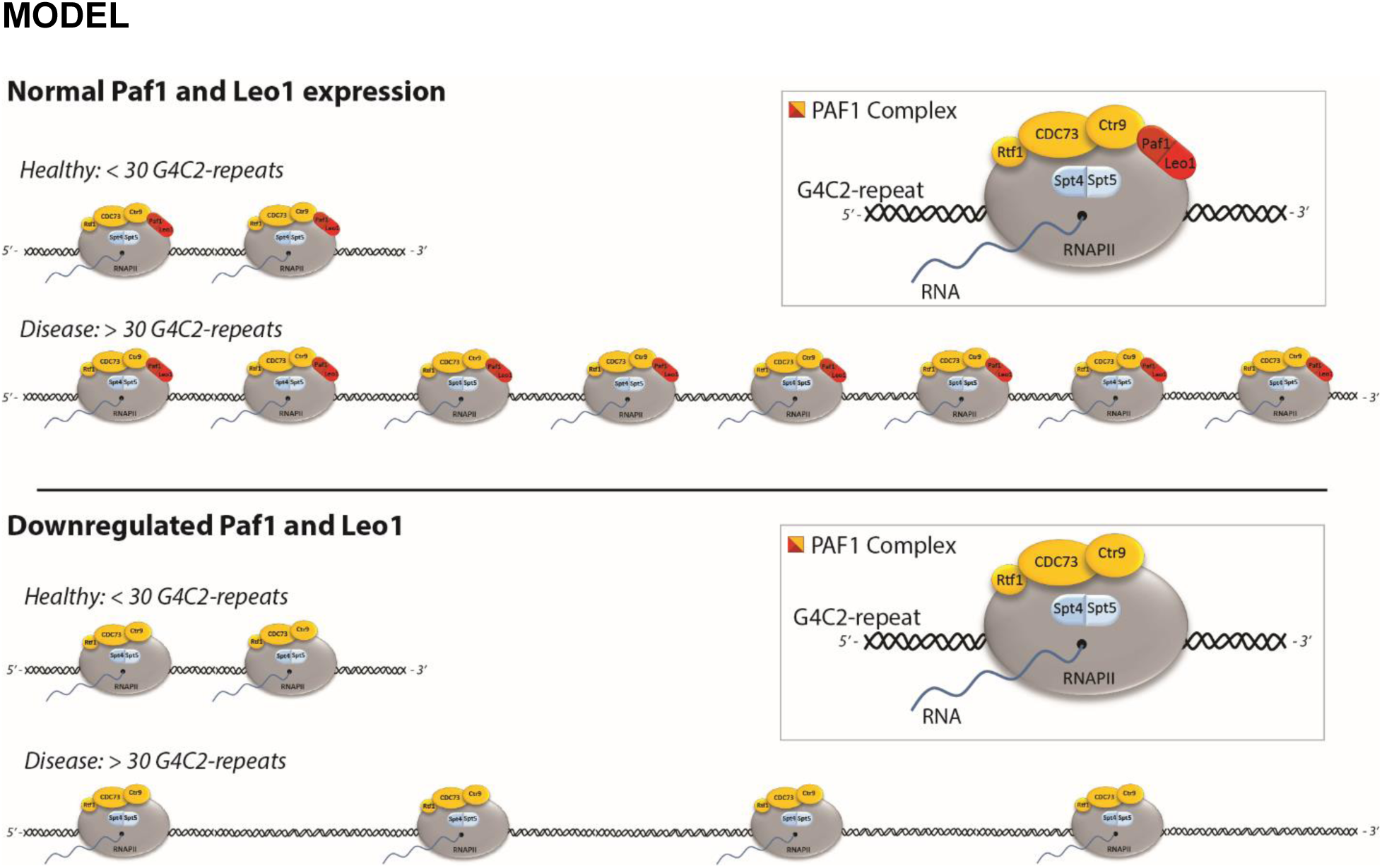

The CDC73/PAF1 complex is recruited to elongating RNAPII at G4C2-repeats. CDC73/PAF1 complex components Paf1 and Leo1 seem most important for RNAPII-driven transcription of expanded (G4C2)n, showing specificity to >30 G4C2 repeats in flies. Further, *PAF1* and *LEO1* are upregulated in response to the expression of >30 G4C2 repeats in both the fly and in C9+ FTD patients due to increased requirements for them. We hypothesize that this is the result of the expanded G4C2 DNA appropriating Paf1/Leo1 along with RNAPII.

## INTRODUCTION

In the *C9orf72* gene, an intronic GGGGCC hexanucleotide (G4C2) expansion of >30 repeats is the most prominent mutation in familial frontotemporal degeneration (FTD) and amyotrophic lateral sclerosis (ALS)^1,2^. These devastating neurodegenerative disorders represent a continuum of the same disease, while it remains unclear how one patient presents with FTD and another with ALS or combined FTD/ALS^3–6^.

While the presence of the expanded G4C2 can confer toxicity through a number of aberrant pathways, amassing evidence supports contributions by gain-of-function mechanisms^7–9^. Accumulation of sense strand G4C2- and antisense strand C4G2-containing RNA results in RNA foci throughout the nervous system and may be toxic^10,11^. Further, these RNAs undergo repeat-associated non-AUG (RAN) translation to produce dipeptide-repeats (DPRs) – GA, GR, GP, PA, and PR – which form potentially toxic aggregates^12,13^. Expressing expanded (G4C2)30+ repeat transcripts, independent of changes to expression of the *C9orf72* gene product, results in toxicity in multiple model systems^7–9^. In *Drosophila*, expression of (G4C2)30+ transgenes results in the formation of RNA foci and expression of DPRs, while causing neurodegenerative effects^14–21^.

Repeat-expansions, including (G4C2)n, have been defined in a number of neurodegenerative diseases^22,23^. Toxicity associated with such mutations can result from aberrant expression of the RNA and/or encoded peptides. As these mutations tend to be GC-rich, they can form secondary structures (e.g. G-quadraplexes) and R-loops which are thought to diminish RNAPII-driven transcription^24–33^. Specialized transcriptional machinery may be required to promote the activity of RNAPII through long repeat expansions. Of note, the DRB-sensitivity-inducing factor (DSIF) complex and the CDC73/PAF1 complex (PAF1C) are RNAPII regulators which may be involved as transcription of GC-rich DNA is sensitive to their loss^34–38^.

DSIF and PAF1C function in RNAPII-driven transcription while they have non-redundant roles in activating RNAPII during elongation^39–42^. DSIF is composed of two proteins, Spt4 and Spt5^40^. Spt4 is implicated as a transcriptional regulator of expanded CAG and G4C2 repeats in disease^20,40,43,44^. PAF1C is composed of five proteins - Paf1, Leo1, CDC73, Ctr9, and Rtf1 and it is unknown if PAF1C has a role in repeat expansion diseases^41,42^. PAF1C downregulation can impact elongation rates of RNAPII, although only a subset of cell-cycle genes are known to require its function for expression^45–50^. In the nervous system, PAF1C is critical to neural differentiation although it remains expressed in mature neurons^51–56^.

Here, we identified the components of PAF1C as modifiers of *C9orf72-associated* disease in an unbiased, large scale RNAi-based screen for modifiers of (G4C2)49-toxicity using *Drosophila*. Downregulation of PAF1C components disrupted transcription of the G4C2 RNA, resulting in reduced toxicity. Importantly, components Paf1 and Leo1 were selective for transcription of long repeat transgenes, unlike Spt4 and the other PAF1C subunits. Furthermore, PAF1C is upregulated in response to the expanded repeat expression in flies as well as in patient tissue. These data highlight the CDC73/PAF1 complex as critical for transcription of the *C9orf72* expanded repeat transcript in disease.

## RESULTS

### RNA Polymerase II transcriptional complexes are enriched among suppressors of (G4C2)49-toxicity

To define cellular mechanisms underlying toxicity that occurs due to expression of expanded (G4C2)30+ repeats *in vivo*, we designed a fly screen utilizing the degenerative eye caused by (G4C2)49 expression with Gmr-GAL4 (Fig. 1a). (G4C2)49-toxicity is seen by disruptions to the external ommatidial organization, red pigmentation, eye size, and internal retinal tissue loss^20^. Using transgenic UAS-RNAi fly lines we defined individual genes whose downregulation altered this toxicity by causing enhancement or suppression^57,58^. Overall, 3,932 genes were tested, ~25% of the fly genome (Supplementary Data). 350 (8.9%) of these altered (G4C2)49-toxicity. These were then rigorously assessed to exclude RNAi lines that caused additive or spurious effects, defined by RNAi that altered the normal eye morphology in control animals or that altered expression of a control UAS-LacZ transgene (Supplementary Fig. 1a-b)^20^. Overall, 119 modifiers of (G4C2)49-toxicity were identified, 55 suppressors and 64 enhancers (Fig. 1b).

**Figure 1:**
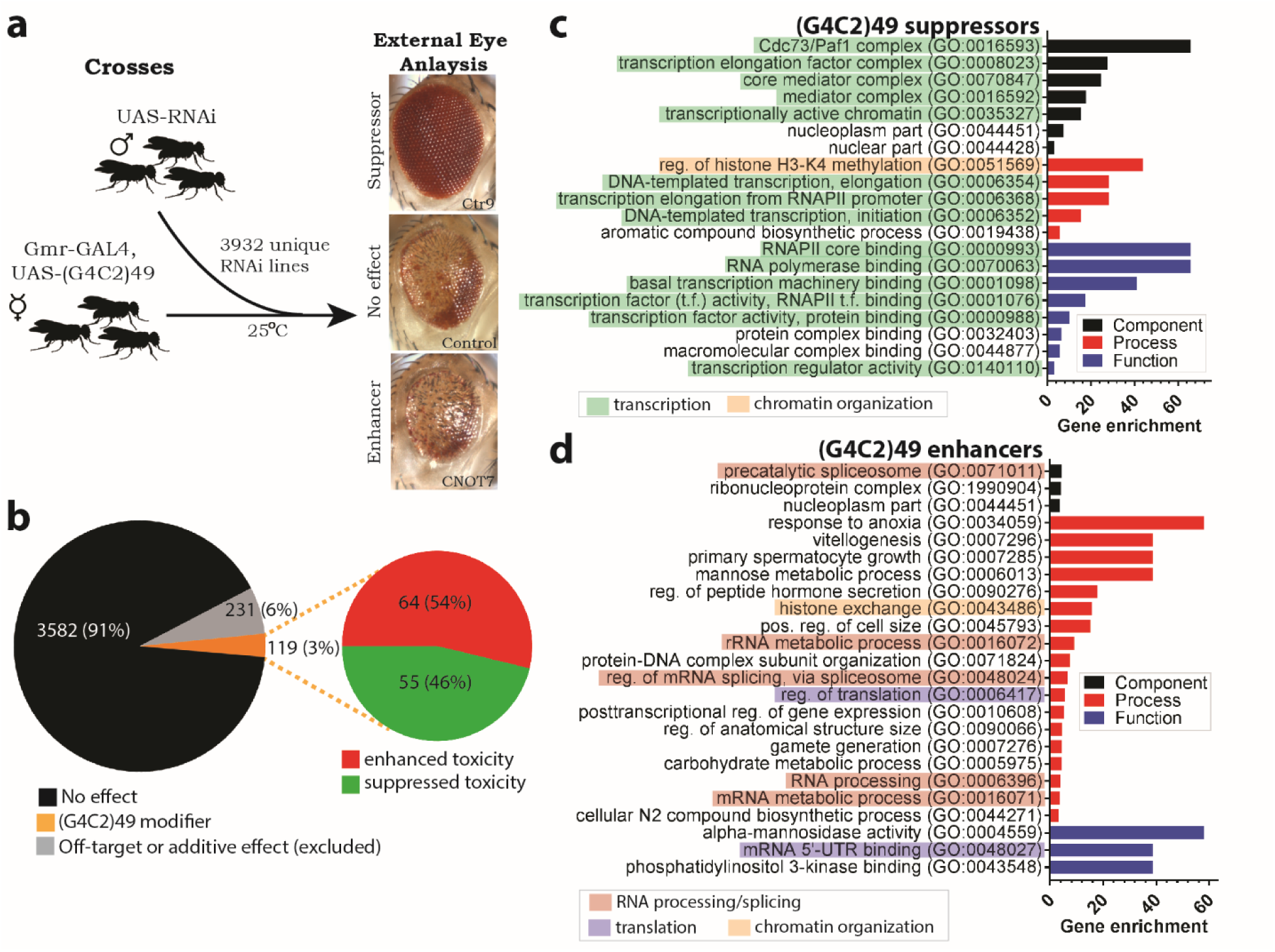
An unbiased genetic screen reveals the CDC73/PAF1 complex as a suppressor of (G4C2)49-induced toxicity in the external fly eye. **(a)** 3,932 RNAi lines were crossed with (G4C2)49-expressing animals. Expression was driven by Gmr-GAL4 (optic system). “Suppressors” reduced degeneration. “Enhancers” increased degeneration. “Lethal enhancers” resulted in no viable adults. Shown are representative images for suppression or enhancement of (G4C2)49-toxicity. **(b)** 119 modifiers of (G4C2)49-toxicity were identified – 55 suppressors and 64 enhancers. Control experiments excluded 231 RNAi lines that had non-specific effects. **(c-d)** Gene ontology analyses revealed GO terms enriched in the panel of suppressors or enhancers found in the screen. Plotted are significant (p-value ≤ 10^^^-3) enrichment scores which represent the degree to which a list of genes in a GO-term are represented within the modifier list^61^.

To determine if these 119 modifiers were enriched for select processes, functions, or components, gene ontology (GO) term analyses were performed^59,60^. Enrichment scores^61^ were plotted for GO-terms that were significantly represented within our panel of modifiers, p-value threshold of 10^^^-3 (Supplementary Fig. 1c; Supplementary Data). Strikingly, among suppressors, there was strong enrichment for genes associated with RNAPII-driven transcription (Fig. 1c; green; Supplementary Data), including components of the CDC73/PAF1 complex (PAF1C) and Mediator complex^62^. Additional transcription regulators included, *ELL* and *Ear* of the Super Elongation Complex (SEC)^63^ and *Spt4* of the DSIF complex. In contrast to suppressors, enhancers of (G4C2)49-toxicity varied in GO-terms. Genes involved in RNA processing and splicing were identified, with components of the pre-catalytic spliceosome being the most enriched complex (Fig. 1d; red; Supplementary Data).

Overall, the PAF1 complex was the most enriched component among transcriptional regulators found to suppress (G4C2)49-toxicity.

### The CDC73/PAF1 complex is selective for toxicity from a G4C2-encoding RNA

Expanded G4C2 RNA produces dipeptide-repeat proteins GR, GA, and GP in the fly^15–18^. Of these, (GR)30+ has been shown to be toxic in the eye of animals^15,21^. As modifiers of (G4C2)49-toxicity may be acting downstream of toxic GR-peptide production, we analyzed the 119 RNAi lines for their ability to modulate (GR)36-toxicity in the fly eye (Fig. 2a; Supplementary Data). The GR fly model used expresses a GR-peptide from a non-G4C2 RNA transcript, thus avoiding potential toxicity caused by a G4C2 repeat-bearing RNA. As the PAF1 and DSIF complexes may be of particular interest in repeat-associated disease^20,34–38,43,44^, RNAi lines targeting components of PAF1C or Spt4 were rigorously examined for modification of (GR)36-toxicity and showed no significant effects (Fig. 2b; Supplementary Fig. 2a-b). Overall, 48 of the 119 modifiers of (G4C2)49 (40.3%) did not similarly alter (GR)36-toxicity, suggesting that they modulate RNA expression or RNA-derived toxicity in the (G4C2)49 animals (Fig. 2b; Supplementary Data)^15,21^. GO-term enrichment of these 48 modifiers revealed that suppression by components of PAF1C was selective to the (G4C2)49 model (Fig. 2c; Supplementary Data). In contrast, the Mediator complex and *ELL* and *Ear* of SEC similarly suppressed both (G4C2)49- and (GR)36-toxicity in the external eye.

**Figure 2:**
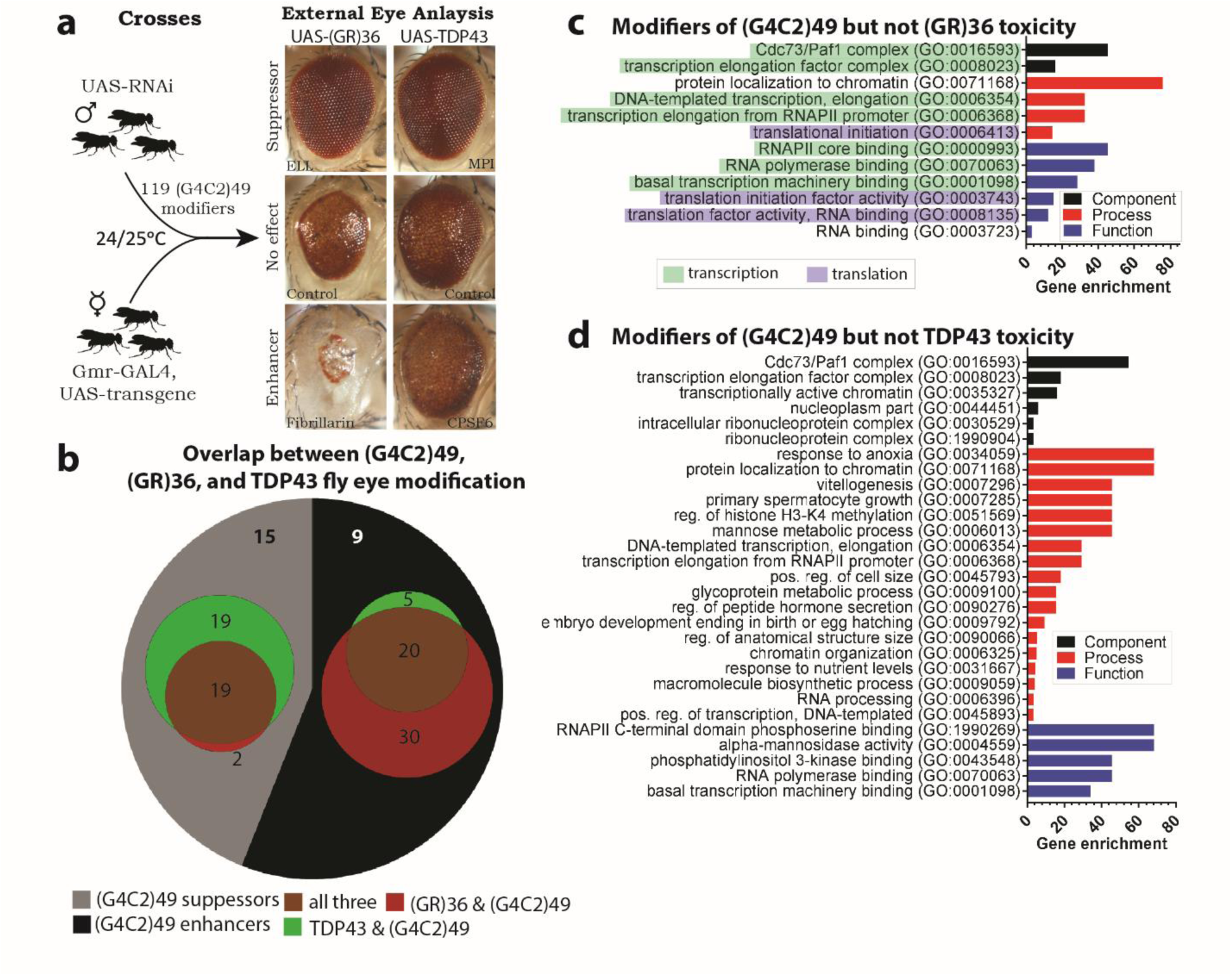
The CDC73/PAF1 complex is not a modifier of (GR)36 or TDP43 *Drosophila* models. **(a)** The 119 (G4C2)49 modifiers were analyzed in (GR)36 and TDP43 fly models to determine if they could act on toxic GR or if they had overlapping effects on TDP43-toxicity. **(b)** Of the 119 modifiers of (G4C2)49 induced toxicity, 71 (59.7%) similarly modified toxicity caused by (GR)36 arguing that these may be acting on the toxic DPR. Alternatively, 63 (52.9%) of the modifiers for (G4C2)49 similarly affect TDP-43 induced toxicity in the eye, arguing overlap between these disease models. **(c)** Gene ontology term analysis of modifiers that *did not* similarly alter (GR)36 toxicity revealed those acting on the (G4C2)49 RNA model. **(d)** Gene ontology term analysis of modifiers that *did not* similarly alter TDP43 toxicity revealed those specific to the expanded G4C2 repeat in C9+ FTD/ALS.

Overall, these data suggest that PAF1C suppression of (G4C2)49-toxicity is the result of effects upstream of toxic-GR production.

### The CDC73/PAF1 complex is selective for the G4C2 expansion

Biological connections between (G4C2)30+ expression and TAR DNA binding protein 43 (TDP43) pathology have been reported in C9+ FTD/ALS patients^64–69^. Thus, we examined the (G4C2)49 modifiers in a TDP43 model to define G4C2-unique pathways (Fig. 2a). Like (G4C2)49, expression of human TDP43 in the fly induces external and internal eye toxicity^70,71^. Rigorous assessment of the RNAi lines targeting PAF1C and Spt4 showed that they had no significant effect on TDP43-toxicity (Fig. 2b; Supplementary Fig. 2c-d). We found that 56 of the 119 (G4C2)49 modifiers (47.1%) were unique to (G4C2)49 (Fig. 2b; Supplementary Data). GO-term analysis revealed that among these, PAF1C was again highly enriched, underscoring specificity of this complex to the expanded G4C2 (Fig. 2d; Supplementary Data). By contrast, other RNAPII-regulators, such as the Mediator complex and *ELL* and *Ear* of SEC, similarly suppressed (G4C2)49- or TDP43-mediated toxicity in the external eye. Furthermore, PAF1C downregulation using RNAi did not alter TDP-43 protein expression (Supplementary Fig. 2e).

Taken together, these data highlight PAF1C selectivity towards (G4C2)49-associated toxicity.

### Downregulation of the CDC73/PAF1 complex suppresses (G4C2)49-induced toxicity in the fly

We further examined features of PAF1C modification in (G4C2)49 expressing flies. We confirmed that the RNAi lines being used to target PAF1C components are effective, causing >80% knockdown of *Paf1, Leo1, CDC73, Ctr9* and 50% knockdown of *Rtf1* (Supplementary Fig. 3a).

Toxicity induced by expanded G4C2 repeat impacts the internal as well as the external eye^20^. When PAF1C components were downregulated in eyes of (G4C2)49 expressing animals, internal retinal tissue loss was reduced (Fig. 3a-b). Notably, *Paf1, Leo1*, and *Rtf1* RNAi resulted in ~80% recovery and *CDC73* and *Ctr9* RNAi caused ~100% recovery as measured by retinal depth (control RNAi = 13.4±6.2μm, Paf1 RNAi = 44.4±4.7μm, Leo1 RNAi = 47.2±10.8μm, CDC73 RNAi = 63±6.1μm, Ctr9 RNAi = 69.1±18.0μm, Rtf1 RNAi = 41.8±12.7μm). This indicates that loss of these components is markedly effective in mitigating toxicity of the expanded G4C2 repeat in the eye. Loss of any of the PAF1C components did not affect the external or internal eye of control animals (Fig. 3c-d).

**Figure 3:**
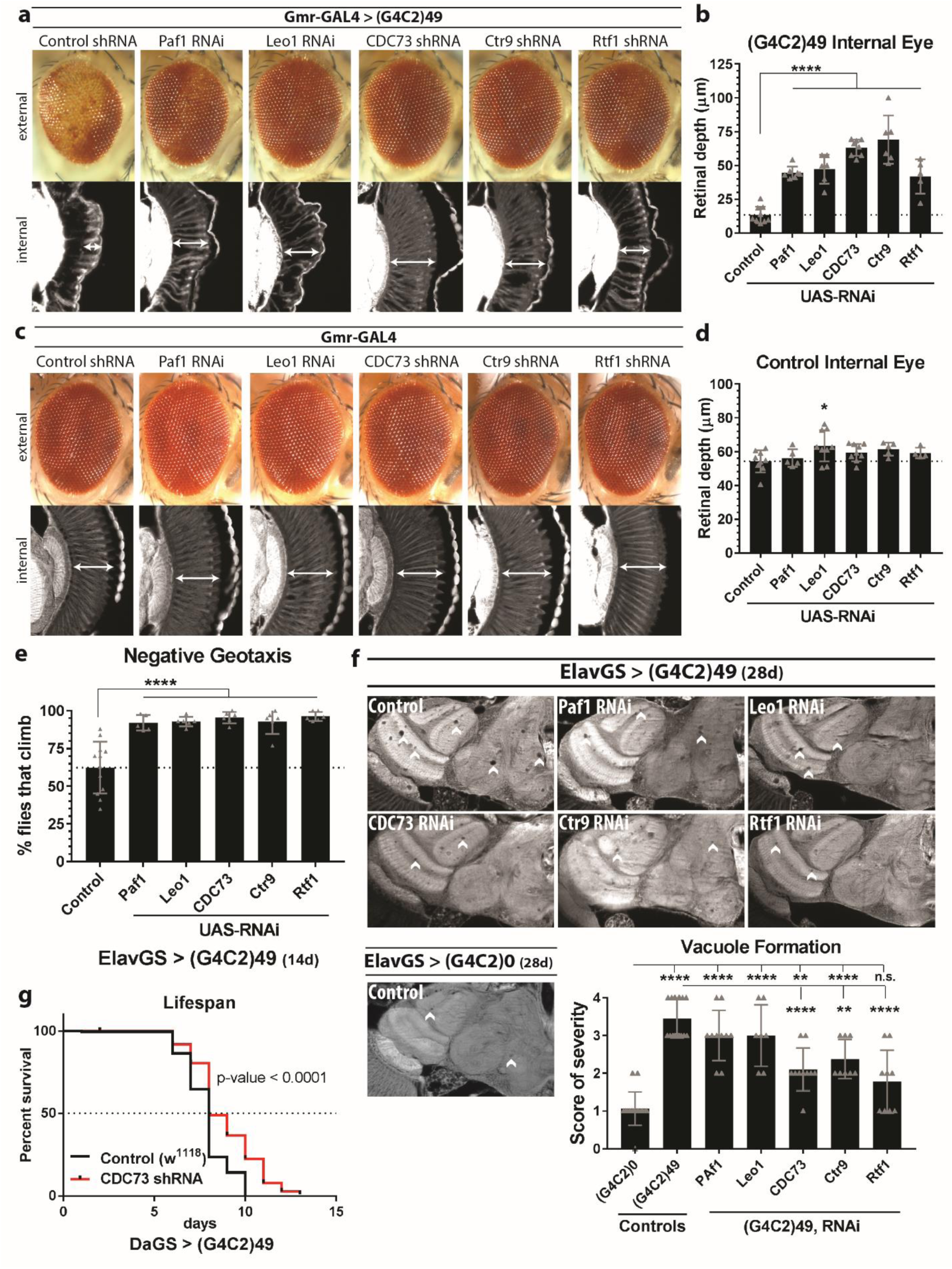
Reduced expression of components of the CDC73/PAF1 complex suppress (G4C2)49-induced toxicity in multiple contexts in the fly. **(a)** Downregulation of each of the components of PAF1C mitigates toxicity associated with (G4C2)49 expression in the fly eye, seen externally by reduced pigment loss and reduced disruption in ommatidial organization and internally by increased integrity of the retinal tissue. **(b)** Quantification of the internal retina depth (n=5-9). **(c-d)** Downregulation of components of PAF1C has no effect on control fly eyes (n=4-10). **(e)** Expression of (G4C2)49 in the adult fly nervous system induces climbing deficits after 14d of expression (ElavGS). Downregulation of PAF1C in these flies rescues climbing deficits (n=98-120). **(f)** Downregulation of PAF1C components mitigate neuron death caused by ElavGS driven expression (G4C2)49 in the brain as seen by vacuole formation (n=7-18). For quantification, a vacuole severity scale was developed where 0 = no vacuoles and 4 = medium/large, frequent vacuoles (Supplementary Fig. 4a). **(g)** Knockdown of CDC73 in adult flies ubiquitously expressing (G4C2)49 results in a lengthened lifespan arguing toxicity induced by (G4C2)49 expression is mitigated (n=197-198). Statistics: ANOVAs for a-f, log-rank test for g, * ≤ 0.05 p-value. All experiments were run multiple times to confirm reproducibility of results. Error bars denote standard deviation.

To examine effects of reduced expression of PAF1C components more broadly in the fly nervous system on (G4C2)49-toxicity, we expressed the (G4C2)49 transgene using a drug-inducible neuronal GAL4 driver, ElavGS. First, we assessed climbing ability of animals. At 14d, only 62% of (G4C2)49 expressing animals were able to climb 4cm up the wall of a vial within 20s (Fig. 3e). Reduced expression of any of the PAF1C components – *Paf1, Leo1, CDC73, Ctr9*, and *Rtf1* –significantly suppressed this effect, such that climbing abilities were maintained at 90-100% (Fig. 3e).

Next, we analyzed animals for age-dependent vacuole formation in the fly brain, an indicator of neurodegeneration^72^. Neuronal expression of (G4C2)49 in adult animals caused large vacuoles throughout the brain by 28d. We found that RNAi lines targeting *CDC73, Ctr9*, and *Rtf1* caused significant reductions in vacuole size and number in the brain of (G4C2)49 expressing animals; mean degeneration score of 3.4±0.5 became reduced to 2.10±0.6 *(CDC73* RNAi), 2.38±0.5 *(Ctr9* RNAi), and 1.78±0.8 *(Rtf1* RNAi), with the normal brain being 1.1±0.4 (Fig. 3f; Supplementary Fig. 4). Further, the strong suppressor *CDC73* could increase survival of these animals, with 50% survival being increased by 24h and the end time point being extended by 72h (Fig. 3g). RNAi targeting other components of the PAF1C showed reduced lifespan in control flies, making it inaccurate to evaluate them as suppressors of (G4C2)49 by lifespan (Supplementary Fig. 3c).

Together, these results support that PAF1C plays an important role in (G4C2)49-induced toxicity in the fly, with downregulation of PAF1C components suppressing toxicity in the eye as well as broadly in the nervous system.

### Paf1 and Leo1 mediate RNA expression from expanded G4C2 in the fly brain

PAF1C has been well characterized for its involvement in RNAPII elongation and is important for transcription of GC-rich DNA^36,41,42^. Given this and the selectivity of downregulated PAF1C to mitigating toxicity in (G4C2)49 expressing animals, we hypothesized that PAF1C may be important for successful transcription of long G4C2 repeats in disease situations.

We first asked if downregulation of PAF1C components – *Paf1, Leo1, CDC73, Ctr9*, and *Rtf1* – affected the amount of RNA produced from expanded (G4C2)49 in the fly nervous system, using *Spt4* RNAi as a positive control (Fig. 4a; red). Downregulation of all PAF1C components caused a significant reduction in RNA produced from the (G4C2)49 transgene – *Paf1* RNAi caused a 40% reduction (SE±0.08), *Leo1* RNAi a 51% reduction (SE±0.04), *CDC73* RNAi a 54% reduction (SE±0.05), *Ctr9* RNAi a 59% reduction (SE±0.06), and *Rtf1* a 60% reduction (SE±0.05). Importantly, these values were similar to *Spt4* RNAi which caused a 49% reduction (SE±0.05) (also^20^).

**Figure 4:**
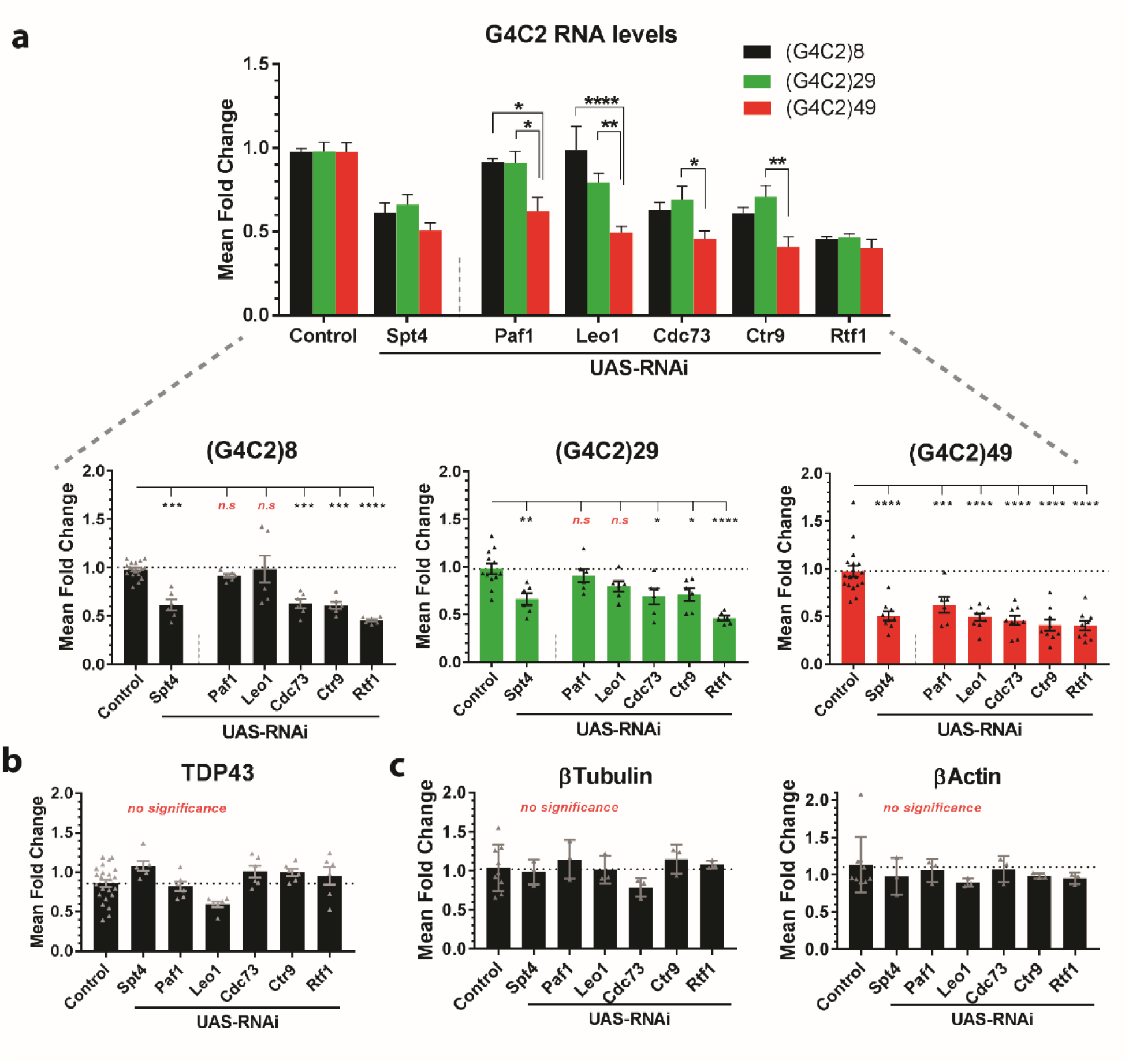
Downregulation of *Paf1* and *Leo1* selectively alters (G4C2)30+ transgene expression in the fly brain. **(a)** (G4C2)8, 29, and 49 transgenes were co-expressed with PAF1C RNAi lines in the adult brain using a drug inducible, neuronal driver (ElavGS, 16d). The level of transgene expression in heads was measured by qPCR. Downregulation of *Paf1* and *Leo1* did not affect RNA levels of (G4C2)8 and (G4C2)29 but caused significant reduction in (G4C2)49 expression. Knockdown of other PAF1C components -- *CDC73, Ctr9*, and *Rtf1* -- and *Spt4* all similarly reduced expression of (G4C2)8, (G4C2)29, and (G4C2)49 transgenes. **(b)** A TDP43 transgene was similarly expressed like (G4C2)n transgenes, using ElavGS (16d, heads) and showed no change in expression by qPCR upon knockdown of PAF1C. **(c)** RNA levels for endogenous RNAPII driven genes, β-Tubulin and β-Actin, are unchanged upon knockdown of PAF1C. The ubiquitous, drug-inducible driver, DaGS, was used to drive RNAi expression in whole animals (6d). Statistics: ANOVAs, * ≤ 0.05 p-value. RNA expression data was normalized to the RNAPI-driven housekeeping gene, RP49. All experiments were run multiple times to confirm reproducibility of results. For a-b, error bars denote standard error. For c, error bars denote standard deviation.

We determined if this effect was specific to the expanded (G4C2)49 transgene, by assessing RNA expression from additional G4C2 repeat-containing transgenes, including a nontoxic (G4C2)8 and an intermediate length (G4C2)29 repeat. Surprisingly, *Paf1* and *Leo1* were selective for expanded (G4C2)49 as *Paf1* and *Leo1* RNAi did not significantly reduce expression from the shorter repeat transgenes (Fig. 4a; black and green respectively). By contrast, *Spt4, CDC73, Ctr9*, and *Rtf1* RNAi caused significant reductions in expression from (G4C2)8 or (G4C2)29 of ≤ 30%, similar to effects on the (G4C2)49 transgene (Fig. 4a; red). To better understand how these numbers compared among the three repeat lengths, data were plotted together allowing for statistical comparisons between each (G4C2)n transcript levels with individual RNAi (Fig. 4a; top). Importantly, *Paf1* or *Leo1* RNAi show no significant difference between the expression level of (G4C2)8 and (G4C2)29 and a statistically significant drop between (G4C2)≤29 and (G4C2)49 RNA levels. By contrast, *Spt4* RNAi and other PAF1C RNAi had statistically similar decreases in RNA levels among short and expanded G4C2 repeat lengths. Together, these findings suggest that, of the components of PAF1C, *Paf1* and *Leo1* are of special importance for expression of longer repeats.

To determine if the reduced expression of G4C2-transgenes caused by PAF1C RNAi was specific to the G4C2 construct, we examined the effects of knockdown on expression of an alternative disease transgene, TDP43. TDP43 transcript levels were not affected in the fly brain (Fig. 4b; Supplementary Fig. 2e). Further, we confirmed that ubiquitous expression of PAF1C did not alter general RNAPII transcription, by examining mRNA levels of endogenous housekeeping genes (Fig. 4c)^20,41,42,45–50^.

Effects of PAF1C RNAi could be the result of co-regulation of the components of the complex – *Paf1, Leo1, CDC73, Ctr9*, and *Rtf1* – with each other and/or *Spt4*. To examine this, we expressed *Paf1, Leo1*, or *CDC73* RNAi in whole animals using the drug-inducible ubiquitous driver, DaGS, and determined whether there was altered expression of the PAF1C components not targeted by the RNAi or *Spt4* by qPCR (Supplementary Fig. 3b). Data supported that loss of PAF1C components does not alter expression of *Spt4* or alternative PAF1C components in adult flies.

Together, these data argue that *Paf1* and *Leo1* are conferring repeat-length specificity to the transcriptional machinery, helping RNAPII transcribe G4C2 repeats longer than 30 units.

### Endogenous CDC73/PAF1 complex is upregulated in response to neuronal expression of expanded G4C2

Given the important role PAF1C is playing in (G4C2)49-expressing animals, we considered that the complex may be dysregulated. To determine this, we examined the expression from endogenous PAF1C genes – *Paf1, Leo1, CDC73, Ctr9*, and *Rtf1 –* in animals expressing the (G4C2)49 repeat in the fly brain using a drug-inducible, neuronal driver, ElavGS. Surprisingly, all components of PAF1C were upregulated (Fig. 5a). Compared to control, the steady state mRNA levels of *Paf1* were increased by 69% (SE±0.10), *Leo1* by 58% (SE±0.13), *CDC73* by 71% (SE±0.17), *Ctr9* by 49% (SE±0.12), and *Rtf1* by 64% (SE±0.05). This upregulation was not seen upon expression of a short (G4C2)8 repeat. Further, PAF1C upregulation appeared selective to the repeat expansion as the transcript levels of PAF1C components were not upregulated in TDP43 expressing animals, but rather downregulated (Fig. 5b).

**Figure 5:**
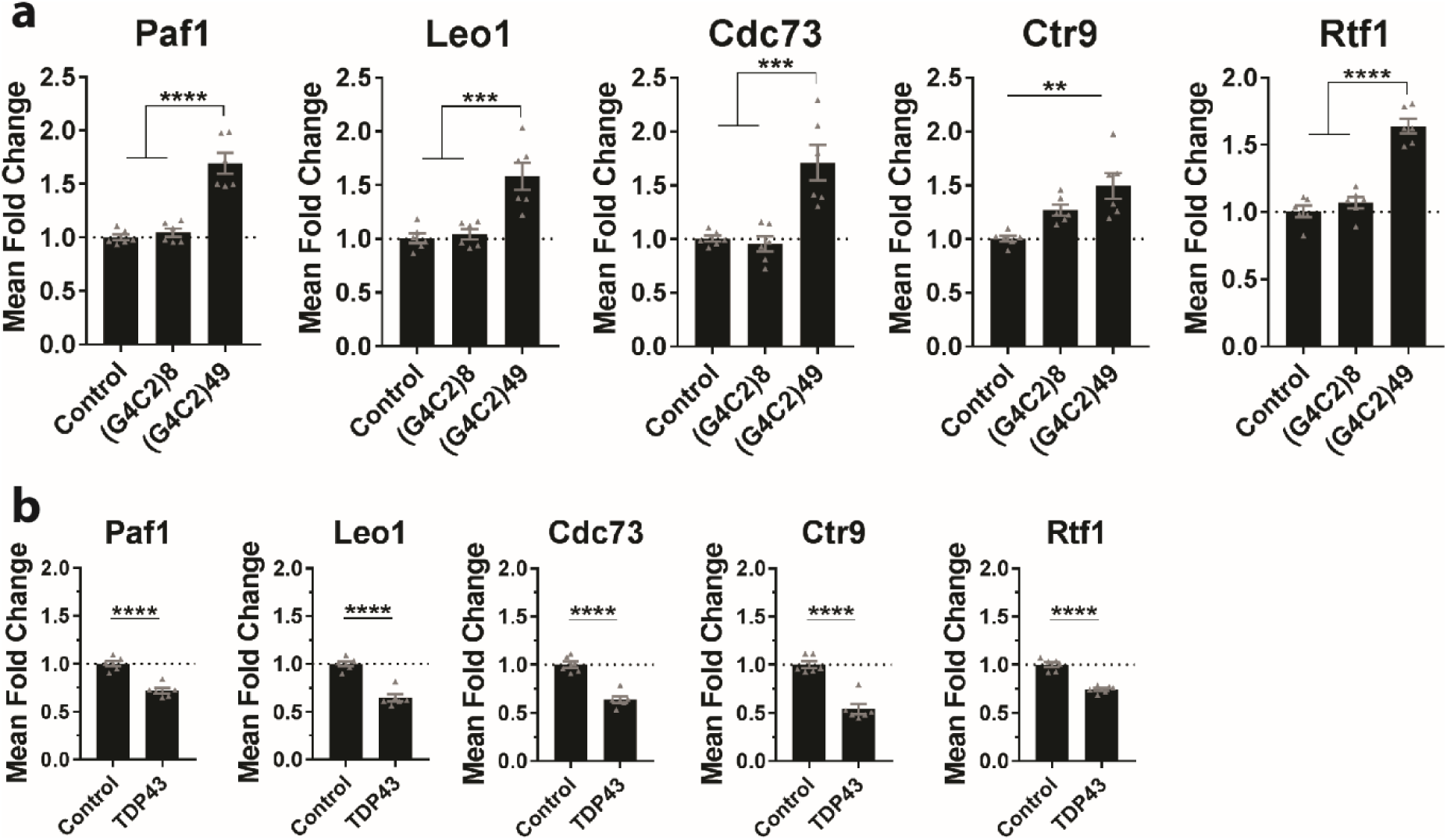
Endogenous CDC73/PAF1 complex is upregulated in response to (G4C2)49 expression in the fly brain. **(a)** RNA was extracted from control, (G4C2)8, and (G4C2)49 expressing animals and analyzed for endogenous expression of PAF1C components by qPCR. Transgenes were expressed in the adult fly nervous system, using ElavGS, while heads were used for analysis (16d). Components of PAF1C are upregulated in (G4C2)49 expressing animals, relative to control or (G4C2)8 expressing animals. Differences in expression are likely underestimated as RNA was extracted from neuronal and non-neuronal tissue in animal heads while transgenes were expressed only in neurons with ElavGS. **(b)** A non-G4C2 disease transgene, TDP43, was similarly expressed and showed no upregulation of PAF1C components. Statistics: ANOVAs in a, student t-test in b, * ≤ 0.05 p-value. RNA expression data was normalized to the RNAPI-driven housekeeping gene, RP49. All experiments were run multiple times to confirm reproducibility of results. Error bars denote standard error.

Taken together, these data indicated that PAF1C becomes upregulated in response to expression of expanded G4C2 repeats in neurons.

### *PAF1* and *LEO1* are upregulated in C9+ FTD and FTD/ALS

The fly data indicated that PAF1C is upregulated in response to expression of an expanded G4C2 repeat in the brain and that PAF1C is required for expression of that repeat. To better understand the role of PAF1C in disease, we extended studies to C9+ FTD, FTD/ALS and ALS samples (patients that have >30 G4C2 repeats in *C9orf72)*. We focused on *PAF1* and *LEO1* as these PAF1C components selectively regulated expression of the longer G4C2 repeat in *Drosophila*.

RNA was extracted from frontal cortex tissue of C9+ patients (n=67), C9-patients (n=56), or healthy controls (n=27) and the level of *PAF1* and *LEO1* expression were assessed by qPCR (Fig. 6a; Supplementary Fig. 5a). In patients diagnosed with FTD, *PAF1* and *LEO1* were significantly upregulated only when the repeat expansion was present – *PAF1* was ~40% upregulated compared to healthy controls and ~30% upregulated compared to C9-FTD; *LEO1* was upregulated ~25% compared to both healthy controls and C9-FTD. Interestingly, in the frontal cortex of ALS cases, neither *PAF1* nor *LEO1* showed upregulation, independent of the presence of the G4C2-repeat expansion. Patients showing symptoms of both FTD and ALS fell between FTD only and ALS only with C9+ FTD/ALS cases showing weaker upregulation of *PAF1* and *LEO1* – *PAF1* was ~30% upregulated and *LEO1* was ~20% upregulated in C9+ cases versus healthy controls.

**Figure 6:**
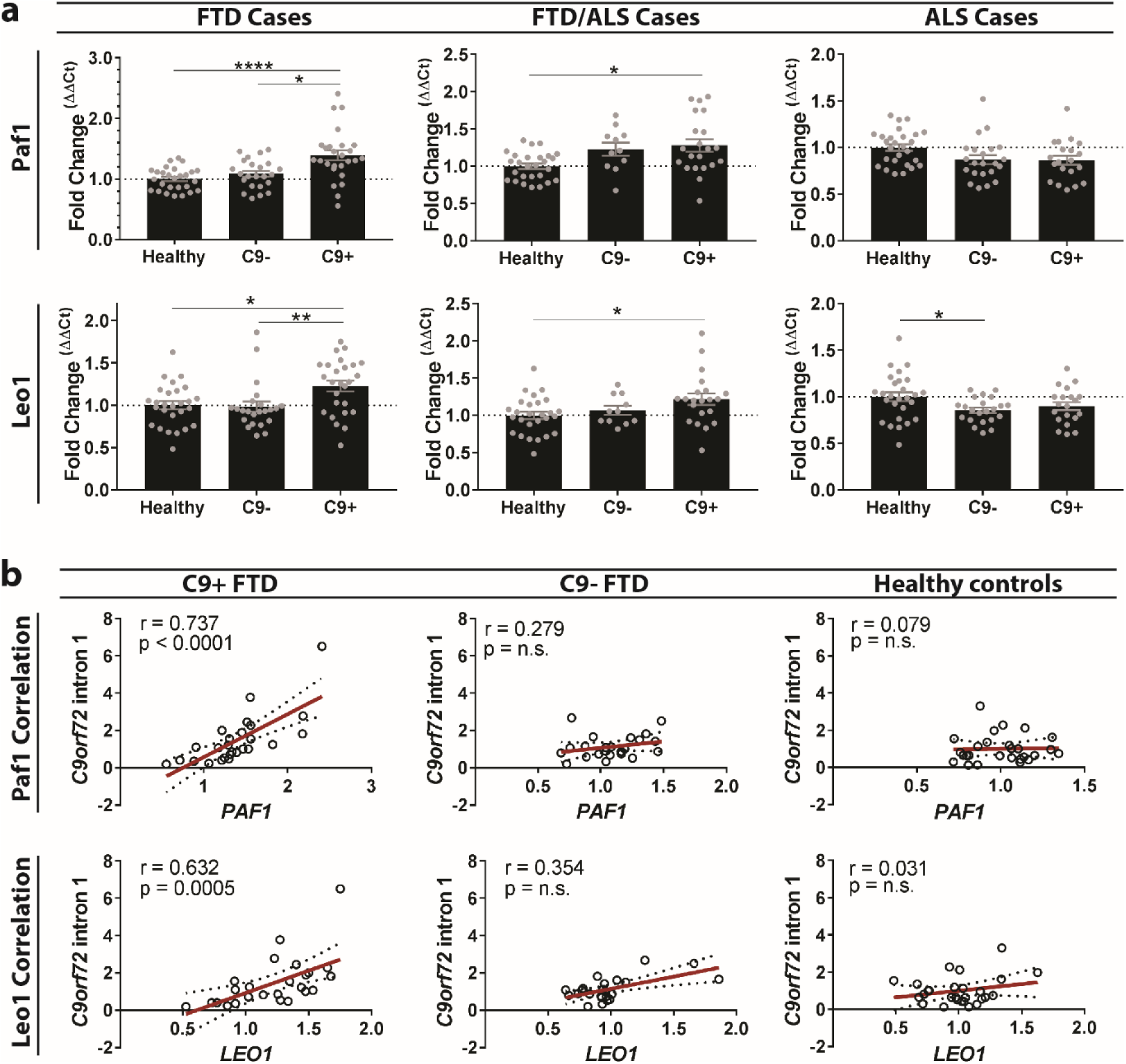
*PAF1* and *LEO1* are upregulated in C9+ FTD and their expression positively correlates to expression of repeat-containing *C9orf72*. **(a)** qPCR analysis of endogenous expression of *PAF1* and *LEO1* in post-mortem, frontal cortex tissue derived from healthy control (n=27), C9-patients (n=56), and C9+ patients (n=67). Data is broken down by diagnosis revealing that in FTD and FTD/ALS cases, *PAF1* and *LEO1* are significantly upregulated, with the largest effects observed in FTD cases. In contrast, frontal cortex tissue from ALS cases do not show an upregulation of *PAF1* and *LEO1*. **(b)** To look for positive correlations in *PAF1* and *LEO1* expression and the expression of repeat-containing *C9orf72*, Spearman correlation coefficients were calculated where 0 means no correlation and 1.0 means 100% correlated. *C9orf72* intron 1: the intronic gene region immediately 3’ of the G4C2 repeat in the C9orf72 pre-mRNA transcript. Strong and significant correlations in expression were observed only in C9+ FTD cases (r > 0.30; p-values < 0.0005). In contrast, C9-FTD and healthy controls show no correlation in expression of the *C9orf72* gene which is lacking the G4C2 expansion. Statistics: * ≤ 0.05 p-value. Error bars denote standard errors.

Altogether, patient data show that *PAF1* and *LEO1* of PAF1C are upregulated in the frontal cortex of C9+ FTD patients but not ALS patients carrying the repeat expansion.

### Expression of *PAF1* and *LEO1* positively correlates to expression of repeat-containing *C9orf72* transcripts in FTD

As *Paf1* and *Leo1* expression were important for the expression of the expanded G4C2 repeat in the fly brain, we considered that *PAF1* and *LEO1* upregulation in C9+ FTD may positively correlate to the expression of *C9orf72* only when the repeat expansion is present. To examine this, Spearman r correlations were performed between the level of *PAF1* or *LEO1* expression and intronic *C9orf72* transcript levels^73,74^. We compared transcripts in C9+ FTD, C9-FTD, and healthy controls. Results showed strong (r > 0.3 to r > 0.6) and significant (p-value <0.0005) positive correlations in *PAF1* and *LEO1* expression with the expression of the repeat-containing *C9orf72* transcript in C9+ FTD cases – for *PAF1* vs *C9orf72*: r = 0.74, 95% CI 0.5 to 0.9; for *LEO1* vs *C9orf72*: r = 0.63, 95% CI 0.3 to 0.8 (Fig. 6b). No correlations were observed in C9-FTD cases or in healthy controls. In C9+ ALS cases, no correlations were observed between *PAF1* or *LEO1* levels compared to the level of repeat-containing *C9orf72* transcript and C9-ALS cases showed a relatively weaker correlation than that observed for C9+ FTD (Supplementary Fig. 5b).

In summary, these data support that *PAF1* and *LEO1* expression positively correlates to the expression of the G4C2 expansion in C9+ FTD. No association was found in frontal cortex of C9+ ALS patients, potentially reflecting the different diagnosis^75–77^.

## DISCUSSION

An unbiased, RNAi-based screen in *Drosophila*, covering ~4000 genes, revealed the CDC73/PAF1 complex (PAF1C) as a *C9or72*-disease modifier. Downregulation of the core components of PAF1C – *Paf1, Leo1, CDC73, Ctr9*, and *Rtf1 –* selectively suppressed (G4C2)49 repeat toxicity in multiple contexts in the fly. Mechanistically, PAF1C suppression was associated with reduced RNA production from the (G4C2)49 transgene in the fly nervous system. Importantly, loss of components *Paf1* and *Leo1*, which form a heterodimer^78^, selectively reduced expression from expanded G4C2 repeats compared to shorter repeats. In contrast, other PAF1C components and *Spt4* showed similar knockdown effects on expression of short and expanded G4C2 transgenes. Moreover, in the fly, PAF1C became upregulated upon expression of (G4C2)49 in neurons. This did not seem to be the result of toxicity or stress as expression of another disease gene, TDP43, did not show increased RNA levels for PAF1C components. Further, extended studies in patients showed that *PAF1* and *LEO1* were upregulated in FTD patients only when they carried the expanded repeat in *C9orf72*. This upregulation positively correlated with expression of the repeat-containing *C9orf72*, supporting a mechanistic link between PAF1C levels, expression of a G4C2-repeat, and C9+ disease.

A total of 119 modifiers of (G4C2)49-toxicity were identified in our RNAi-based screen (Fig 1; Supplementary Data). Remarkably, gene-ontology (GO) term analysis of the 55 suppressors showed significant enrichment for RNA polymerase II (RNAPII) transcriptional regulators – including PAF1C, the Mediator complex, *ELL* and *Ear* of the Super Elongation Complex (SEC), and *Spt4* of the DSIF complex. Although we focused on PAF1C, the other RNAPII regulators may also be important in disease, potentially relating to unique transcriptome changes associated with C9+ situations^66,73,79^. Among the 64 enhancers of (G4C2)49-toxicity identified, RNA processing and splicing factors were prominent. Curiously, the majority of these enhancers similarly modulate (GR)36-toxicity (Fig. 2). These data are consistent with reports of RNA dysregulation and splicing deficits in C9+ FTD/ALS, while suggesting that disruptions in RNA metabolism may result from toxic GR^80^.

RNAPII-driven transcription across GC-rich DNA is hypothesized to be difficult due to the propensity of the DNA to form secondary structures, such as R-loops and G-quadraplexes^24–33^. Given this, the activity of multiple elongation factors may be required for efficient transcription through expansions like (G4C2)30+. *Spt4* was previously implicated as a transcriptional regulator of CAG and G4C2 repeat expansions in disease^20,43,44^. Our data indicates that PAF1C is also critical for mediating expression of G4C2 repeats in FTD/ALS (Fig. 4). *Paf1* and *Leo1* of PAF1C seem particularly important for RNAPII-transcription of expanded G4C2 as loss of these two components selectively results in reduced expression of a (G4C2)49 transgene in the fly (Fig. 4). Consistent with their having similar effects on (G4C2)49 expression, these two components form a heterodimer that is important for PAF1C activation of elongating RNAPII^37,49,81,78,82–84^.

In contrast to Paf1/Leo1, Spt4 showed similar effects on different G4C2 repeat lengths, an unexpected result given previous findings (Fig. 4)^20,43^. Differences in how Spt4 and Paf1/Leo1 loss impact expanded G4C2 expression may be due to their distinct roles during transcription^39–42^. Evidence suggests that Spt4 primarily acts during the transition of RNAPII from poised to elongation or during transcription termination^38,49,85–87^. In contrast, Paf1/Leo1 seems to act primarily during elongation^46,49,82–84^. Compellingly, Spt4 and Paf1C are reported to interact. In yeast, Spt4 recruits PAF1C via the Rtf1 subunit during the transition of RNAPII from initiation to elongation^85,88,89^. However, in higher organisms, including *Drosophila*, the dependence of PAF1C on Spt4 for recruitment to elongating RNAPII is less clear. Rtf1 is less tightly associated with other PAF1C components while recent work shows that PAF1C recruitment can be Spt4-independent in mammals^46,48,90–92^. Further, PAF1C directly interacts with elongating RNAPII^81^. Together, data suggests that PAF1C is playing a unique role in transcription elongation across a G4C2-repeat expansion.

FTD and ALS represent a spectrum of the same disease while overlap may be the result of unique genetic backgrounds of individuals^4–6,75–77^. To date, only Ataxin-2 has been suggested as a disease modifier underlying the different diagnoses^3^. We see upregulation of *PAF1* and *LEO1* in the frontal cortex of C9+ FTD, but not C9+ ALS cases compared to C9-cases and healthy controls (Fig. 6). Importantly, the frontal cortex is thought to be a primary brain region resulting in FTD-associated symptoms^75,76,93^. Further, C9+ FTD/ALS cases have lower levels of upregulation of *PAF1* and *LEO1* than C9+ FTD cases, supporting that these would fall between FTD only and ALS only disease. Patient data were consistent with fly data, as flies expressing (G4C2)49 showed upregulation of endogenous PAF1C components (Fig. 5). Overall, it is tempting to hypothesize that modulation of *Paf1* or *Leo1* may be among a number of mechanisms contributing to the onset of different disease phenotypes in C9orf72-associated disease.

PAF1C may be an attractive therapeutic candidate for C9+ FTD. In yeast, *PAF1C* has been reported to be non-essential^20,41,43^. In the fly, *Leo1* has also been shown to be non-essential^94^. In mice, *PAF1^+/−^* or *LEO1^+/−^* heterozygosity yields no obvious abnormalities, although *PAF1^−/−^* or *LEO1^−/−^* null animals show pre-weaning lethality^95–97^. Other components of PAF1C follow this same trend, with heterozygous mice showing no or few effects^95–97^. While Spt4 (non-essential in yeast) was previously proposed as a potential therapeutic target in C9+ FTD/ALS^20^, it may have more critical organismal functions than PAF1C in higher organisms; *SUPT4A^+/−^* (murine *Spt4*) heterozygous mice are viable but show a number of abnormalities and *SUPT4A^−/−^* null mice are embryonic lethal^95–97^. Altogether, we hypothesize that specific components of PAF1C, like *Leo1*, may represent a potential therapeutic target for *C9orf*72-associated disease^45,46,48,49^.

This study presents the first evidence that PAF1C is an important player in *C9orf72-associated* disease, particularly in C9+ FTD. Further investigations into PAF1C may define its impact in other neurodegenerative diseases that result from aberrant expression of repeat expansions^22,23^.

**Author contributions**
This work was done by LDG under the mentorship of NMB. MP performed patient studies and analyses under the mentorship of LP. LFRM added technical support during PAF1C-focused studies, including qPCRs and lifespans. ML added technical support during screening and performed initial control GAL4/UAS-LacZ westerns post-screening. MP performed paraffin sectioning for internal eye and vacuole formation investigations.

## Acknowledgments

We thank Amit Berson, Jason Kennerdell, Leeanne McGurk, Janani Saikumar, Ananth Srinivasan and other members of the Bonini laboratory for helpful comments. Decklan P. Cerza and Alexander Chen provided minimal technical support. We thank the Transgenic RNAi Project (TRiP) at Harvard Medical School (NIH/NIGMS R01-GM084947) and the Vienna *Drosophila* Research Center for developing transgenic RNAi fly stocks used in this study. This work was funded by the Systems and Integrative Biology NIH training grant (T32-GM07517), NIH R01-NS078283, NIH R35-NS09727, and ALS Association.

